# Individual kinetochore-fibers locally dissipate force to maintain robust mammalian spindle structure

**DOI:** 10.1101/846154

**Authors:** Alexandra F. Long, Pooja Suresh, Sophie Dumont

**Author notes:** Corresponding authors: Sophie Dumont, Alexandra Long.

## Abstract

At cell division, the mammalian kinetochore binds many spindle microtubules that make up the kinetochore-fiber. To segregate chromosomes, the kinetochore-fiber must be dynamic and generate and respond to force. Yet, how it remodels under force remains poorly understood. Kinetochore-fibers cannot be reconstituted *in vitro*, and exerting controlled forces *in vivo* remains challenging. Here, we use microneedles to pull on mammalian kinetochore-fibers and probe how sustained force regulates their dynamics and structure. We show that force lengthens kinetochore-fibers by persistently favoring plus-end polymerization, not by increasing polymerization rate. We demonstrate that force suppresses depolymerization at both plus- and minus-ends, rather than sliding microtubules within the kinetochore-fiber. Finally, we observe that kinetochore-fibers break but do not detach from kinetochores or poles. Together, this work suggests an engineering principle for spindle structural homeostasis: different physical mechanisms of local force dissipation by the k-fiber limit force transmission to preserve robust spindle structure. These findings may inform how other dynamic, force-generating cellular machines achieve mechanical robustness.

## Introduction

The spindle segregates chromosomes at cell division, and must do so accurately and robustly for proper cell and tissue function. In mammalian spindles, bundles of 15-25 microtubules called kinetochore-fibers (k-fibers) span from the kinetochore at their plus-ends to the spindle pole at their minus-ends (Rieder, 1981; McDonald et al., 1992; McEwen et al., 1997). K-fibers are dynamic at both ends (Mitchison, 1989; Cassimeris and Salmon, 1991), and we now have a wealth of information on the molecular regulation of their dynamics (Cheeseman and Desai, 2008; Bakhoum and Compton, 2012; Monda and Cheeseman, 2018). To move chromosomes, k-fibers generate force through plus-end depolymerization (Koshland et al., 1988; Grishchuk et al., 2005; Mitchison et al., 1986). Yet, while we are beginning to understand how the mammalian k-fiber generates force (Inoué and Salmon, 1995; Grishchuk, 2017), we know much less about how force from the k-fiber and surrounding spindle in turn affects k-fiber structure and dynamics. Defining this relationship between k-fibers and their mechanical environment is central to understanding spindle structural homeostasis and function.

Force affects microtubule dynamics and structure in a variety of contexts (Dogterom et al., 2005). From *in vitro* experiments coupling single microtubules to yeast kinetochore particles, we know that force can regulate all four parameters of microtubule dynamic instability (Akiyoshi et al., 2010; Sarangapani et al., 2013): it increases polymerization rates while slowing depolymerization, and favors rescue over catastrophe. From *in vivo* experiments, we know that force exerted by the cell correlates with changes in mammalian k-fiber dynamics (Wan et al., 2012; Dumont et al., 2012; Auckland et al., 2017), and that reducing and increasing force can bias k-fiber dynamics in different systems (Nicklas and Staehly, 1967; Skibbens et al., 1995; Khodjakov and Rieder, 1996; Skibbens and Salmon, 1997; Long et al., 2017). However, the feedback between force, structure and dynamics in the mammalian k-fiber remains poorly understood. For example, we do not know which dynamic instability parameters are regulated by force, or at which microtubule end. Similarly, we do not know how microtubules within the k-fiber remodel their structure (e.g. slide or break) under force, or the physical limits of the connections between k-fibers and the spindle. These questions are at the heart of understanding how the spindle can maintain its structure given its dynamic, force-generating parts (Oriola et al., 2018; Elting et al., 2018). Addressing these questions requires the ability to apply force on k-fibers with spatial and temporal control, while concurrently imaging their dynamics. Yet, exerting controlled forces in dividing mammalians cells remains a challenge, and mammalian spindles and k-fibers cannot currently be reconstituted *in vitro*. Chemical and genetic perturbations can change forces on k-fibers *in vivo,* but these alter microtubule structure or dynamics, either directly or indirectly through regulatory proteins (De Brabander et al., 1986; Vladimirou et al., 2013; Alushin et al., 2014). Thus, direct mechanical approaches are needed inside mammalian cells.

Here, we use glass microneedles to directly exert force on individual k-fibers inside mammalian cells and determine how their structure and dynamics remodel under sustained force. Inspired by experiments in insect spermatocytes (Nicklas and Staehly, 1967; Nicklas, 1997; Lin et al., 2018), we sought to adapt microneedle manipulation to pull on k-fibers in mitotic mammalian cells for many minutes while monitoring their dynamics with fluorescence imaging. We show that forces applied for minutes regulate k-fiber dynamics at both ends, causing k-fiber lengthening, but do not cause sliding of the microtubules within them. Further, we demonstrate that sustained forces can break k-fibers rather than detach them from kinetochores or poles. Thus, k-fibers respond as a coordinated mechanical unit – remodeling at different sites to locally dissipate force, while preserving the connections between chromosomes and the spindle. Together, these findings suggest local force dissipation as an engineering principle for the dynamic spindle to maintain its structure and function under force and for other cellular machines to do the same.

## Results

### Microneedle manipulation of mammalian spindles enables sustained force application on k-fibers with spatial and temporal control

To determine how mammalian k-fibers remodel under force, we sought an approach to apply forces with spatial and temporal control for sustained periods, compatible with cell health and live imaging of structure and dynamics. We adapted microneedle manipulation to pull on individual k-fibers in mammalian cells (Fig. 1A) and developed methods to do so gently enough to exert force for several minutes (Suresh et al., 2019). We used PtK cells as these are large and flat, have few chromosomes which allows us to pull on individual k-fibers, and are molecularly tractable (Udy et al., 2015). We used a micromanipulator and a fluorescently labeled glass microneedle to contact a target metaphase PtK cell. We used microneedles with a diameter of 1.2 ± 0.1 µm in the z-plane of the k-fiber. Pulling on an outermost k-fiber in the spindle for several minutes, we could reproducibly exert controlled forces, moving the microneedle with specific velocities over any given duration (Fig.1B) and direction. The microneedle only locally deformed the cell membrane and spindle and remained outside of the cell, allowing precise, local control of where force is applied (Fig. 1C) (Suresh et al., 2019). Upon careful removal of the microneedle, cells typically entered anaphase (Fig. 1D). These observations are consistent with cell health maintenance during these sustained manipulations. Thus, we can use microneedle manipulation to exert forces with spatial and temporal control over minutes on a mammalian k-fiber, and thereby probe how force regulates k-fiber structure and dynamics.

**Figure 1.**
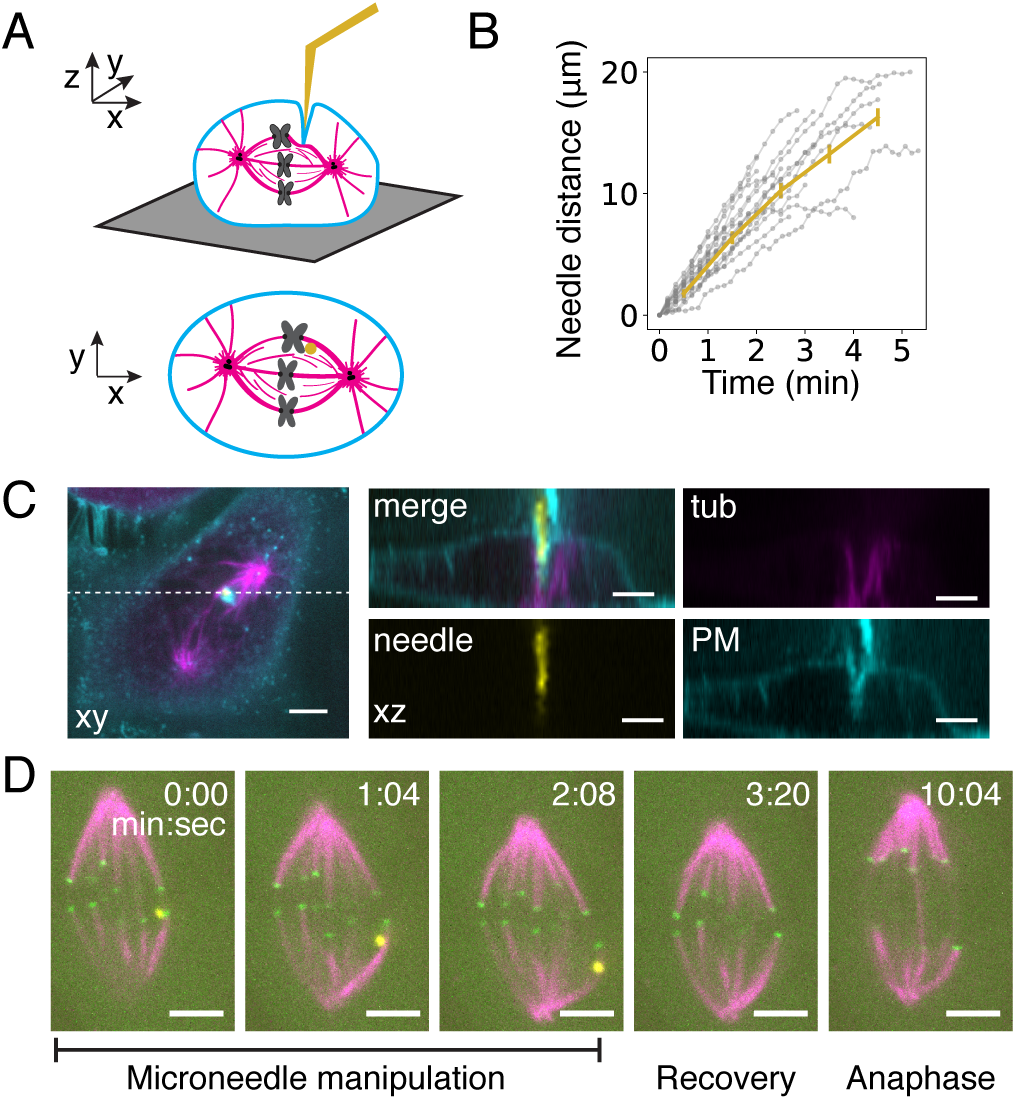
Microneedle manipulation of mammalian spindles enables sustained force application on k-fibers with spatial and temporal control. **A)** Cartoon representation of microneedle (yellow) placement (3D and cross-section) in a metaphase mammalian cell to exert sustained force on a k-fiber. **B)** Plot of linear microneedle displacement over time during manipulation in metaphase PtK cell (mean ± SEM, n = 18 cells). This approach allows smooth, reproducible pulls on single mammalian k-fibers. **C)** Representative z-stack reconstruction shows geometry of microneedle contact with the cell and metaphase spindle (GFP-tubulin, magenta) as diagrammed in (**A**). The plasma membrane (CellMask Orange dye, cyan) locally deforms around the microneedle (Alexa-647, yellow) but does not alter whole cell shape or puncture the cell. Scale bar = 4 µm. **D)** Representative timelapse images of microneedle (Alexa-555, yellow) manipulation to exert force on a k-fiber: it displaces the metaphase spindle (Cdc20-YFP, green; SiR-tubulin, magenta) and deforms the pulled k-fiber. Manipulated spindles typically progress to anaphase (here at 10:04). Scale bar = 4 µm. See also Video 1.

### Individual mammalian k-fibers switch to persistent lengthening in response to sustained applied force

To probe the response of k-fibers to force, we placed the microneedle along the k-fiber, within a few microns of the outermost sister kinetochore pair (Fig. 2A,B). We moved the microneedle at a speed of 5.2 ± 0.2 µm/min for 3.1 ± 0.3 min (Fig. 1B), perpendicular to the spindle’s long axis at the start of manipulation. We predicted that in response to force from the microneedle the spindle would either locally or globally deform (Fig. 2A). In response to this perturbation, the spindle translated and rotated, with faster microneedle speeds giving rise to faster spindle speeds (Fig. 2C,D). Thus we see global movement of the spindle in response to force. Yet, in these same spindles we also observed that k-fibers lengthened, indicating that the spindle also locally responds to force (Fig. 2E). During the pull, the manipulated k-fiber bent and lengthened by 4.1 ± 0.8 µm; meanwhile, an unmanipulated k-fiber in the same spindle half lengthened significantly less over the same duration (net k-fiber growth 0.03 ± 0.32 µm, Mann-Whitney U test, p = 6×10^−5^, Fig. 2F). Thus, force is dissipated locally by k-fiber bending and lengthening, and globally by whole spindle movements.

**Figure 2.**
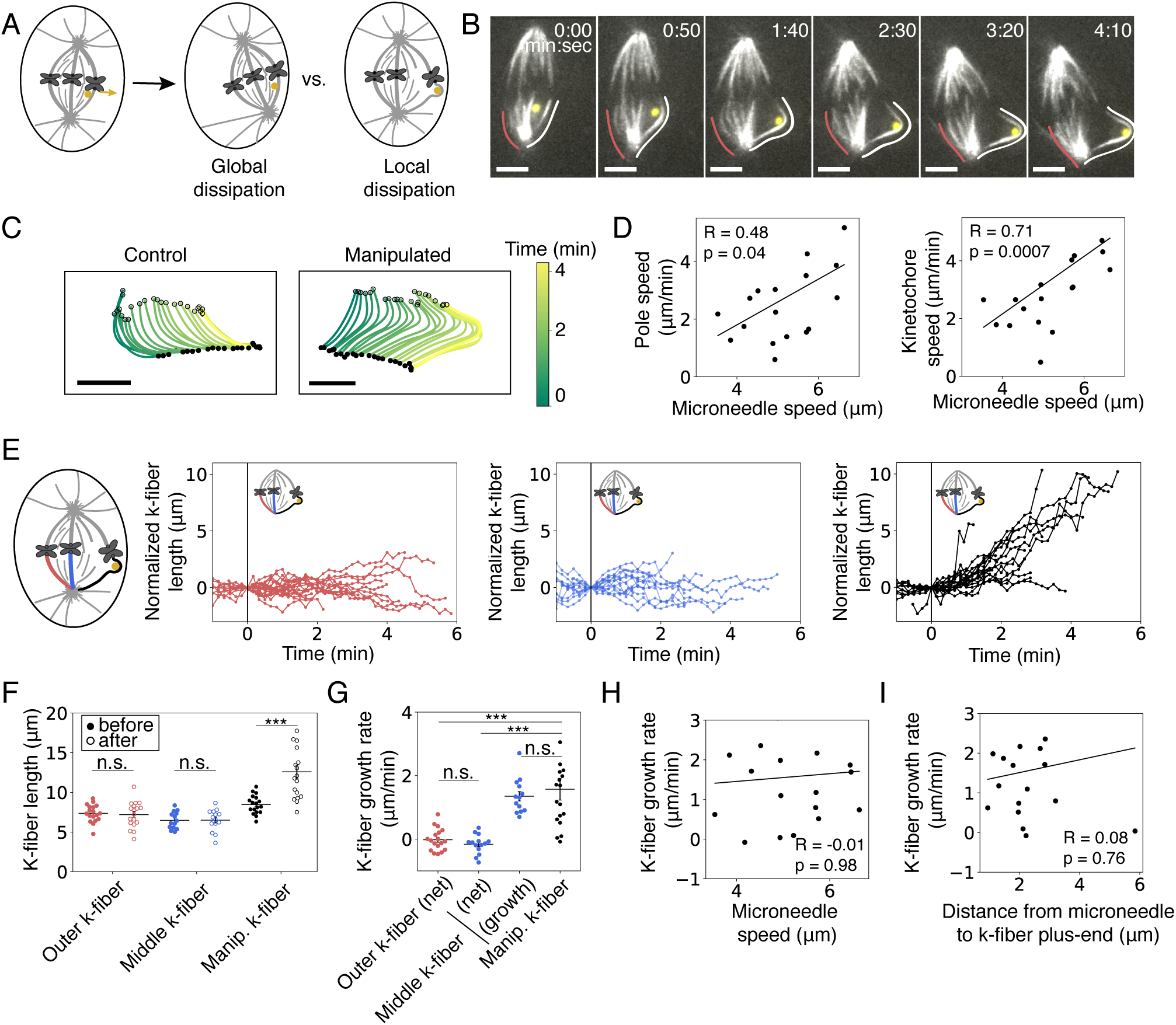
Individual mammalian k-fibers switch to persistent lengthening in response to sustained applied force. **A)** Assay to locally exert force on an outer k-fiber using a microneedle (yellow circle) to probe its response to force (yellow arrow). Possible outcomes include global movement of the whole spindle and local deformation of the k-fiber, reflecting global and local dissipation of applied force, respectively. **B)** Representative timelapse images of spindle and k-fiber (SiR-tubulin, white) movement and remodeling in response to sustained force from a microneedle (Alexa-555, yellow) as in Fig. 1B. The whole spindle rotates and translates while the k-fiber proximal to the microneedle (white line, tracked) bends and lengthens compared to a control k-fiber (red line, tracked). Scale bar = 4 µm. See also Video 2. **C)** Maps of the tracked k-fiber shapes and positions for control and manipulated k-fibers from (**B**). Open circles indicate plus-end positions and filled circles indicate pole positions. The manipulated k-fiber (right) translates in the XY plane and bends and lengthens over time; the control k-fiber (left) similarly translates but does not lengthen. **D)** Speed of proximal pole (left) and plus-end (kinetochore, right) movement relative to the speed of microneedle movement within a half-spindle. Half-spindle movement is positively correlated with microneedle speed, indicating global dissipation of force (pole: Spearman R = 0.48, p = 0.04; plus-end: Spearman R = 0.72, p = 0.0007, n = 18 cells). **E)** K-fiber length as a function of time, normalized by subtracting the initial length at start of force application (t = 0) for k-fibers manipulated (right, black, n = 18 cells), in the middle of the half-spindle (middle, blue, n = 13 cells), and on the opposite side of the half-spindle (left, red, n = 18 cells). The micromanipulated k-fiber lengthens persistently during force application while the other k-fibers grow and shrink but don’t systematically change length. **F)** Average k-fiber lengths at start and end of force application as a function of k-fiber position in the half-spindle. The manipulated k-fiber (black, n = 18 cells) significantly increased in length (p = 0.0002, Wilcoxon signed-rank test) while the middle and outer k-fiber lengths remain unchanged (p = 0.73, n = 13 cells and p = 0.35, n = 18 cells, Wilcoxon signed-rank test). Plot shows mean ± SEM. **G)** Plot of average k-fiber growth rate for manipulated k-fibers (black, n = 18 cells) compared to middle k-fibers (blue, n = 14 cells) or outer k-fibers (red, n = 18 cells) in the same half-spindle. Only the manipulated k-fiber lengthened significantly during force application while neighboring k-fibers continued oscillating between lengthening and shortening phases (manipulated k-fiber versus middle k-fiber ‘net’, p = 1.6×10^−5^, manipulated k-fiber versus outer k-fiber ‘net’, p = 1.4e05, middle k-fiber ‘net’ compared to outer k-fiber, (p = 0.3, Mann-Whitney U test). The growth rate of the manipulated k-fiber was not significantly different than the growth rate of the middle k-fiber during just the growth phases of its oscillations (blue ‘growth’, p = 0.98, Mann-Whitney U test). Plot shows mean ± SEM. **H)** Growth rate of the manipulated k-fiber as a function of the speed of microneedle movement. The growth rate of the manipulated k-fiber did not correlate with the speed of microneedle movement (Spearman R = 0.21, p = 0.46, n = 18 cells). **I)** Growth rate of the manipulated k-fiber as a function of distance between the microneedle center and the k-fiber plus-end. The growth rate of the manipulated k-fiber does not correlate with the proximity of the microneedle to the plus-end (Spearman R = 0.04, p = 0.88, n = 18 cells).

The manipulated k-fiber grew at 1.6 ± 0.3 µm/min, which was not significantly faster than its neighboring unmanipulated k-fiber during the growth phases of its oscillations (1.4 ± 0.1 µm/min, Mann-Whitney U test, p = 0.98, Fig. 2G). However, the manipulated k-fiber persistently lengthened (Fig. 2E), with either undetectable or very transient shortening, for longer than typical metaphase oscillations (Wan et al., 2012; Civelekoglu-Scholey et al., 2013). There was no correlation between k-fiber growth rate and pulling speed (Fig. 2H), suggesting either that force was dissipated before reaching the k-fiber’s ends or that force does not regulate its maximum growth rate (Nicklas, 1983, 1988; Skibbens and Salmon, 1997; Betterton and McIntosh, 2013). Further, the k-fiber growth rate did not vary with the proximity of the microneedle to the plus-end (Spearman R coefficient = 0.08, p = 0.76, Fig. 2I), which we hypothesized would lead to more direct force transmission, consistent with force not regulating the k-fiber’s maximum growth rate. Together, these findings indicate that individual k-fibers remodel under sustained force for minutes by persistently lengthening. They also suggest that force inhibits their normal switching dynamics rather than substantially increasing their growth rate, which may serve as a protective mechanism to limit the rate of spindle deformations and thereby preserve spindle structure.

### Force on individual mammalian k-fibers suppresses depolymerization at both ends without altering plus-end polymerization rates or inducing microtubule sliding

Metaphase mammalian k-fibers typically depolymerize at their minus-ends, and switch between polymerizing and depolymerizing at their plus-ends. Thus, force could lengthen k-fibers by increasing plus-end polymerization rates, by suppressing depolymerization at either end, by sliding microtubules within the k-fiber (Fig. 3A), or by a combination of these. To determine the physical mechanism of k-fiber lengthening under sustained force, we photomarked PA-GFP-tubulin on a k-fiber before micromanipulation and tracked the photomark’s position and size within the k-fiber (co-labeled with SiR-tubulin) (Fig. 3B) over time. In unmanipulated cells, photomarks fluxed towards the pole at a constant rate that reports on depolymerization at the minus-end (Fig. 3C) (Mitchison, 1989). Upon external force from the microneedle, the photomark to pole distance remained constant (Fig. 3D), while the photomark to plus-end distance increased (Fig. 3E). This response indicates that applied force suppresses microtubule depolymerization at k-fiber minus-ends and that k-fibers lengthen by sustained polymerization at plus-ends.

**Figure 3.**
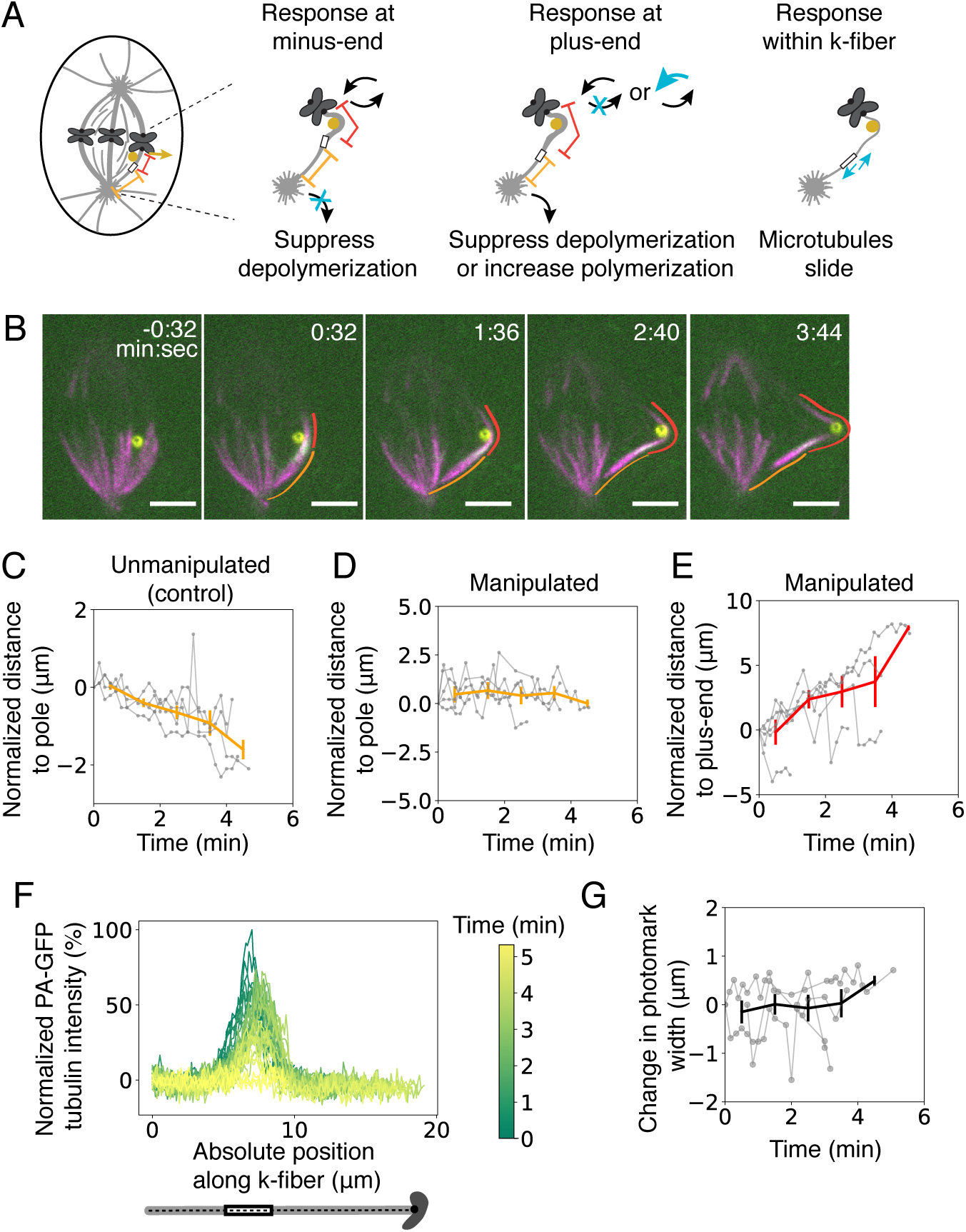
Force on individual mammalian k-fibers suppresses depolymerization at both ends without altering plus-end polymerization rates or inducing microtubule sliding. **A)** Assay to determine the physical mechanism of k-fiber lengthening under force by tracking position of a photomark on the k-fiber during microneedle manipulation. Possible outcomes are shown, not mutually exclusive: the photomark could remain fixed relative to the pole indicating a suppression of minus-end depolymerization (left, blue ‘X’), the position of the photomark to the kinetochore could increase continuously, indicating a suppression of plus-end depolymerization or increase in plus-end polymerization rate (middle, blue ‘X’ or arrow), or the photomark could remain in a fixed position but widen, indicating sliding of microtubules within the k-fiber (right, blue ‘X’). **B)** Representative timelapse images of photomark (PA-GFP tubulin, white) during microneedle (Alexa-555, yellow) manipulation of a k-fiber (SiR-tubulin, magenta). The distance between the photomark and the pole remains constant (orange line) while the distance between the photomark and plus-end increases (red line). Scale bar = 4 µm. See also Video 3. **C)** Plot of the photomark to the pole distance change over time due to flux of microtubules in unmanipulated cells, as a baseline (n = 4 cells). **D)** Plot of the photomark to pole distance change during microneedle manipulation, showing that photomark movement poleward due to microtubule depolymerization is suppressed (n = 4 cells). **E)** Plot of the photomark to plus-end position distance change during microneedle manipulation, showing that k-fibers persistently polymerize at their plus-ends under force (n = 4 cells). **F)** Representative example of photomark intensity linescans over time during manipulation, from same cell as (**B**). **G)** Change in full-width at half-max photomark intensity at each timepoint during microneedle manipulation, showing that photomarks do not widen under force, and thus that there is no detectable microtubule sliding within the k-fiber (n = 4 cells).

Mapping these findings to the previous experiment measuring k-fiber lengthening (Fig. 2E,G), in the subset of k-fibers that lengthened (15/18), the growth rate was 1.9 ± 0.4 µm/min, which is the rate of plus-end polymerization given that depolymerization at both ends is inhibited (Fig. 3D,E). This is similar to the plus-end polymerization rate of neighboring unmanipulated k-fibers during natural growth: lengthening at 1.4 ± 0.1 µm/min (Fig. 2G) while depolymerizing at the minus-end at ∼ 0.5 µm/min results in a polymerization rate of ∼1.9 µm/min at plus-ends (Mann-Whitney U test, p = 0.55) (Long et al., 2017). This indicates that the applied force does not increase mammalian k-fiber plus-end polymerization rates.

Notably, the average width of the photomark remained constant during manipulation (Fig. 3F,G), indicating the microtubules do not detectably slide within the bundle. Thus, the k-fiber behaves as a single coordinated mechanical unit, rather than as microtubules that independently respond to force. Together, our findings indicate that individual k-fibers lengthen under force by remodeling their ends, and not their bundle structure: force suppresses depolymerization locally at both plus- and minus-ends (Fig. 3), leads to persistent plus-end polymerization at a force-independent rate (Fig. 2,3), and does so with the k-fiber responding as a single mechanical unit (Fig. 3). Thus, force is dissipated locally at k-fiber ends. This may limit force transmission to the rest of the spindle, thereby preserving overall k-fiber and spindle architecture for proper chromosome segregation.

### The interfaces between mammalian k-fibers and the kinetochore and pole are more robust than k-fiber bundles under sustained force

Finally, we asked how k-fiber structure and spindle connections changed over the ∼5-7 min lifetime of its microtubules (Gorbsky and Borisy, 1989; Cassimeris et al., 1990; Zhai et al., 1995), since this could set a timescale for their response to force. We hypothesized that as microtubules turn over the manipulated k-fiber could, for example, detach from the spindle or break (Fig. 4A). We used microneedles to pull on k-fibers for several minutes. Over these sustained pulls, we never observed k-fiber detachment from the kinetochore or pole, indicating strong anchorage at those force-dissipating sites (Nicklas and Staehly, 1967; Begg and Ellis, 1979; Nicklas et al., 1982; Gatlin et al., 2010; Fong et al., 2017). Instead, k-fibers bent, lengthened, and then occasionally broke, 3.7 ± 0.5 min after the start of pulling (Fig. 4B). To probe the mechanism of this breakage, we examined k-fiber structure over time and the kinetics of breakage. K-fibers that broke sustained high curvature for many minutes before breaking (Fig. 4C), and reached a maximum curvature similar to those that did not (p = 0.25 Mann-Whitney U test, Fig. 4D). Further, k-fiber breakage kinetics appeared independent of the specific manner in which forces are exerted on the k-fiber: the time to breakage was similar when we moved the microneedle for a shorter time and held it in place, or pulled continuously for the entire duration of manipulation (Fig. 1B, 4E). Together, these suggested that the breakage process occurred gradually over sustained force, rather than rapidly by reaching an acute mechanical limit of k-fiber bending (Nicklas et al., 1989; Gittes et al., 1993; Ward et al., 2014; Schaedel et al., 2015). A k-fiber damage process that is gradual would promote breakage in response to sustained but not transient forces, setting a limiting timescale for restoring spindle structural homeostasis.

**Figure 4.**
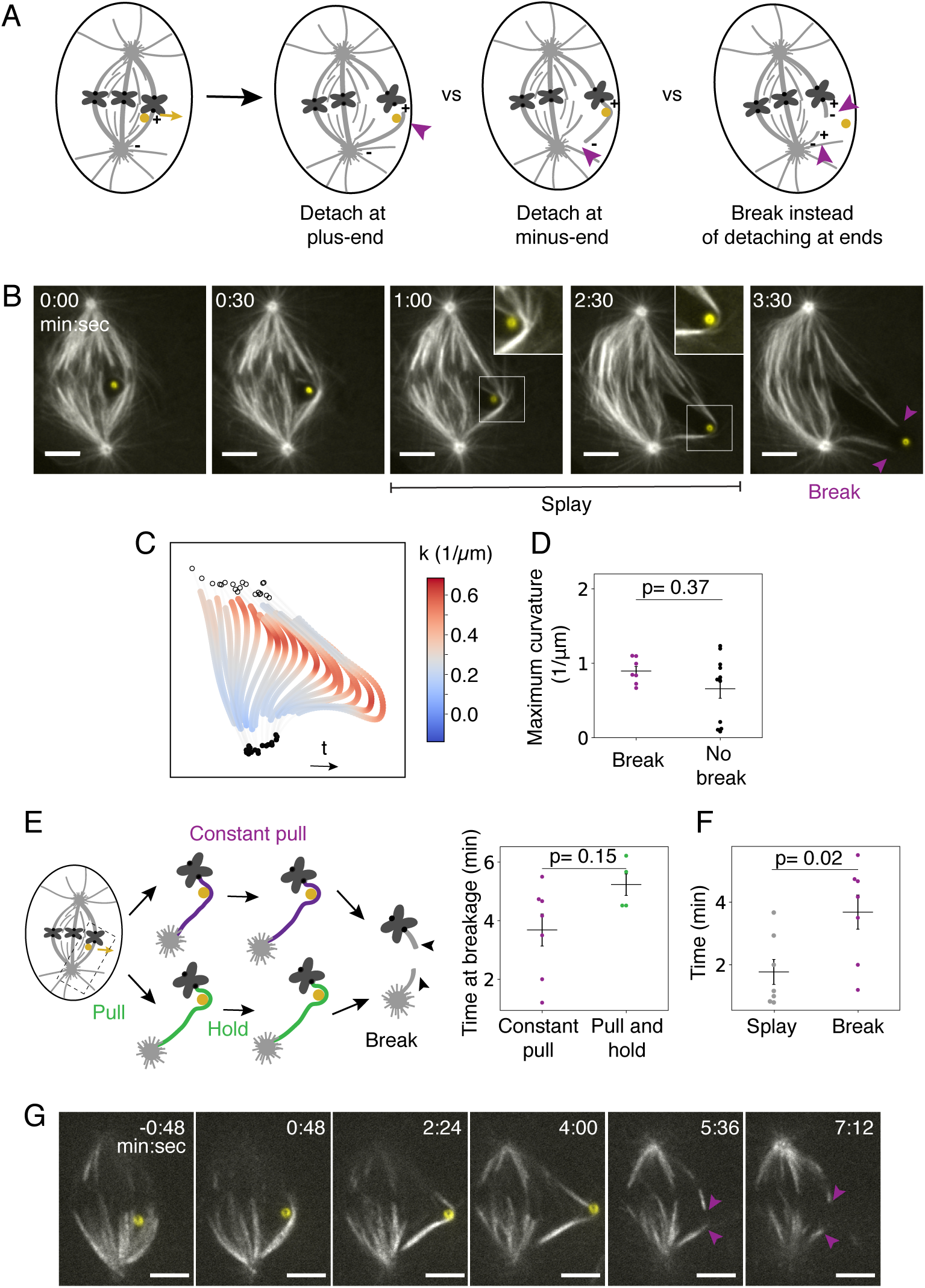
The interfaces between mammalian k-fibers and the kinetochore and pole are more robust than k-fiber bundles under sustained force. **A)** Assay to probe how the k-fiber response to sustained force for minutes. Three example outcomes of force application (yellow arrow) are shown: the k-fiber could detach (purple arrow) from the kinetochore (left), the k-fiber could detach (purple arrow) from the pole (middle), or the k-fiber could remain attached at its ends but break (purple arrows) in its center (right). **B)** Representative timelapse images of k-fiber (GFP-tubulin, white) bending, lengthening and breaking under sustained force. Before the k-fiber breaks, microtubules appear (insets) on the outside of the deformed k-fiber near the area of high curvature next to the microneedle (Alexa-647, yellow). The break creates new microtubule bundle plus-ends (purple arrowheads). Scale bar = 4 µm. See also Video 4. **C)** Example map of local curvature (*k*) along a k-fiber bundle during sustained microneedle manipulation. As the k-fiber bends over time, high curvature (dark red) increases near the microneedle and persists for many minutes before breakage occurs (3.5 min). Open circles indicate plus-end positions and filled circles indicate pole positions. **D)** Maximum curvature along the k-fiber in the last tracked timepoint before breakage in cells with breakage events (purple, n = 6 cells) or at the end of the manipulation for cells with no breakage (black, n = 11 cells, plot shows mean ± SEM, p = 0.37, Mann-Whitney U test). **E)** Cartoon of two different micromanipulation assays that lead to k-fiber breakage: (top, purple) microneedle is moved continuously at 5.2 ± 0.2 µm/min for 3.1 ± 0.3 minutes, (bottom, green) microneedle is moved at 4.5 ± 0.7 µm/min for 1.7 ± 0.2 min and then held in place until breakage. Plot showing no significant difference in the time at breakage in each assay (plot shows mean ± SEM, n = 7 cells and 4 cells, p = 0.15, Mann-Whitney U test). **F)** Plot of the average time to a splaying event (where newly visible microtubules appear near the area of high curvature) and average time to breakage for the subset of cells in which both events occurred. Splaying events occurred significantly before breakage events (plot shows mean ± SEM, n = 9 cells, p = 0.007, Wilcoxon signed-rank test). **G)** Example timelapse images of breakage event in which the newly created bundle plus-ends (lower purple arrow) are highly stable and persist for minutes after breakage. This example cell is the same as shown in Fig. 3B but here displaying the full response including breakage. See also Video 5.

A possible model for gradual damage of the k-fiber over minutes is loss of microtubules as they turn over and fail to replenish within the k-fiber. In addition to turnover, it is also possible that there are alterations to k-fiber microtubule structure that would lead to gradual damage. During these manipulations, we observe microtubule plus-ends that appear to ‘splay’ from the bundle near the needle in 80% of k-fibers before breakage (Fig 4B,F), and when we can track plus-ends after breakage, they fail to depolymerize (Fig 4G). This is in contrast to abruptly created k-fiber plus-ends which depolymerize within seconds (Spurck et al., 1990; Sikirzhytski et al., 2014; Elting et al., 2014) and suggests a change in local microtubule structure prior to breakage that stabilizes plus-ends at the breakage site (Schaedel et al., 2015; Portran et al., 2017; Vemu et al., 2018; McNally and Roll-Mecak, 2018; Gasic and Mitchison, 2019). Together, these findings show how mammalian k-fibers gradually respond to and dissipate sustained forces over their microtubule’s lifetime. They robustly remain attached at kinetochores, yet eventually they locally break in the middle of the bundle, thereby preserving connections of chromosomes to the spindle at the expense of non-essential direct connections to poles (Sikirzhytski et al., 2014; Elting et al., 2014).

## Discussion

In mammals, chromosome segregation is powered by dynamic k-fibers that both generate and respond to force. Here, we use microneedle manipulation to directly probe how k-fiber dynamics and structure respond to sustained force (Fig. 1). We thereby define how the spindle’s longest-lived microtubule structure (Gorbsky and Borisy, 1989; Cassimeris et al., 1990; Zhai et al., 1995) remodels under force, which is key for understanding spindle structural homeostasis. We find that individual k-fibers respond to and dissipate sustained force by locally turning off microtubule depolymerization at both plus- and minus-ends (Fig. 2, 3), and eventually breaking on the timescale of their microtubule turnover (Fig. 4). They do so without increasing their rate of plus-end polymerization (Fig. 2,3), without sliding their microtubules within the k-fiber (Fig. 2,3) and without detaching from kinetochores or poles (Fig. 4). Thus, how the k-fiber responds – and doesn’t respond – to force allows it to act as a single mechanical unit that can maintain its connections to chromosomes and preserve global spindle structure. The ability to directly exert force on the mammalian spindle is key to this work as it allowed us to clearly probe the feedback between force, structure, and dynamics in the spindle (Elting et al., 2018). Together, these findings suggest different physical mechanisms of local force dissipation as an engineering principle for the spindle to maintain its structure and function under sustained forces (Fig. 5). More broadly, this study provides a framework for understanding how the spindle remodels under force during chromosome segregation.

**Figure 5.**
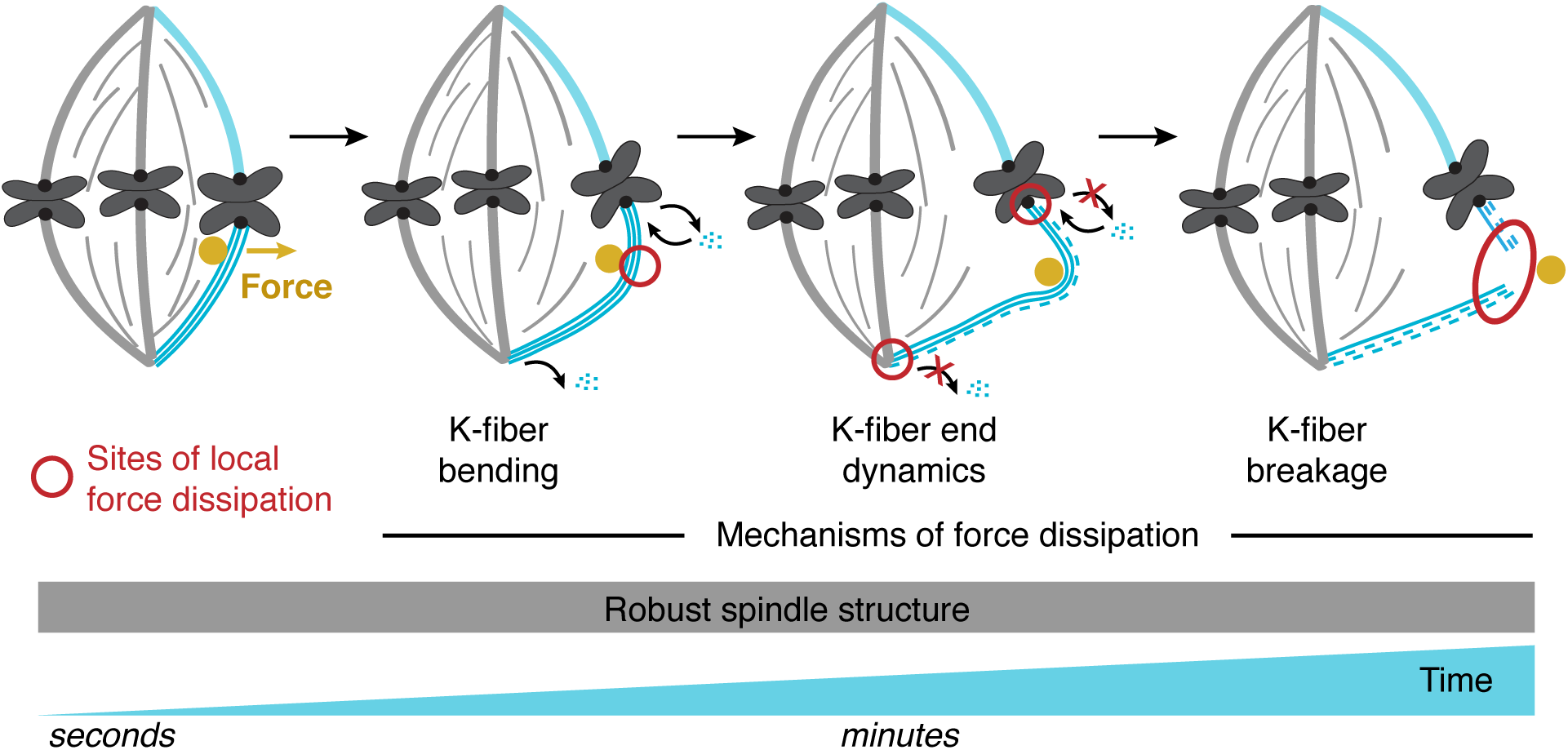
A model for local force dissipation by individual k-fibers to maintain robust mammalian spindle structure. Using micromanipulation to apply sustained forces (yellow circle, arrow) on individual mammalian k-fibers reveals that they locally dissipate force (red circles) using different physical mechanisms over different timescales (blue ramp, dashed lines indicate microtubule turnover) to robustly preserve global spindle structure (gray box). Key to this model is how k-fibers both remodel under and resist sustained force. K-fibers *remodel* and locally dissipate force: they bend (second panel), lengthen through suppressing depolymerization at their plus- and minus-ends (third panel, small black ‘off’ arrows with red ‘X’), and gradually break (fourth panel). In turn, k-fibers also *resist* force to preserve spindle structure: they do not increase their polymerization rate (small black ‘on’ arrows), slide their microtubules, or detach from kinetochores or poles under force. Note that for simplicity, we do not diagram whole spindle movements and only show individual microtubules for the manipulated k-fiber. Thus, local dissipation and isolation mechanisms together preserve mammalian spindle structure under sustained forces: the former limit how far and for how long forces can be transmitted across the spindle, while the latter limit the spindle’s deformation rate and preserve k-fiber and spindle structure and their connections. Together, this model suggests local force dissipation as an engineering principle for the dynamic spindle and other cellular machines to robustly maintain their structure and function under force.

We show that mammalian k-fiber plus-ends persistently polymerize at normal rates in response to applied force (Fig. 2,3). In contrast, microtubules attached to yeast kinetochore particles *in vitro* polymerize faster at higher force, in addition to suppressing catastrophe and favoring rescue under force (Franck et al., 2007; Akiyoshi et al., 2010). In newt cells, force induces persistent k-fiber lengthening at normal k-fiber growth rates (Skibbens and Salmon, 1997), and our findings suggest that this may occur through regulation of dynamics at both ends. The different force-velocity relationships at kinetochore-microtubule plus-ends in mammals and yeast kinetochore particles could, for example, stem from differences in applied forces, kinetochore architecture (Long et al., 2019), or additional regulation in cells. The molecular basis of potential “governors” of k-fiber plus-end polymerization velocity has been a long standing question (Nicklas, 1983; Betterton and McIntosh, 2013; Long et al., 2017), and our findings suggest that in mammals this molecular “governor” is not mechanically regulated. Notably, force not regulating mammalian k-fiber polymerization velocity (Fig. 2,3) could provide a protective upper limit to how fast the spindle can remodel. It also has implications for mechanical communication in the spindle, for example how force regulates kinetochore-microtubule attachments (Li and Nicklas, 1995; Sarangapani and Asbury, 2014).

We demonstrate that force not only regulates the dynamics of individual k-fibers’ plus-ends, but also of their minus-ends (Fig. 3). Thus, both k-fiber ends serve as sites of force dissipation, allowing forces exerted on k-fibers to be locally and robustly dissipated, thereby limiting disruption to the rest of the spindle. The fact that force regulates minus-end dynamics of single k-fibers indicates that their regulation occurs at the level of the individual k-fiber, and not globally as hypothesized when force was applied to the whole spindle (Dumont and Mitchison, 2009; Guild et al., 2017). Though we cannot exclude it, we did not detect force-induced polymerization at k-fiber minus-ends, and thus force dissipation also appears limited at minus-ends. The microneedle approach we present here, combined with perturbations of microtubule regulators at minus-ends (Ganem et al., 2005; Ganem and Compton, 2006), will be key in defining the molecular basis of the regulation of k-fiber minus-end dynamics by force. Together, the response of individual k-fibers’ dynamics to force, at both ends, allows each k-fiber to locally isolate and dissipate applied force while retaining its internal organization and global spindle structure. Therefore k-fiber end dynamics mechanically buffer global spindle deformations from local forces to maintain structural homeostasis (Maddox et al., 2003; Matos et al., 2009).

On longer timescales, we find that the k-fiber breaks under force, without detaching from the kinetochore or pole (Fig. 4). This is surprising as force-induced detachments from kinetochores occur *in vitro* (Akiyoshi et al., 2010) and in meiotic insect cells (Nicklas, 1967; Nicklas and Koch, 1969; Paliulis and Nicklas, 2004; Lin et al., 2018). This difference could, for example, arise from variations in force application, or in the physical properties or architectures of their kinetochores (Cheerambathur et al., 2017; Auckland et al., 2017; Agarwal et al., 2018; Yoo et al., 2018). Instead of detaching, the k-fiber breaks on a timescale similar to that of its microtubule lifetime, suggesting that the k-fiber’s lifetime may limit the long-term impact force can have over spindle structure. Our findings suggest a model of gradual k-fiber damage, and that sustained force may not only regulate biochemistry at the k-fiber’s ends, but also in its middle along the microtubule lattice (Fig. 4F,G). Local defects in the lattice can replenish GTP-tubulin, creating stable sites for microtubule repair or enzymatic activities that may alter the physical properties of microtubules (Schaedel et al., 2015; Portran et al., 2017; Vemu et al., 2018; McNally and Roll-Mecak, 2018; Gasic and Mitchison, 2019). Under sustained force, k-fiber attachments to chromosomes are prioritized over direct connections between chromosomes and poles, which are not necessary for segregation (Elting et al., 2014; Sikirzhytski et al., 2014) and thus may not be key for function.

Altogether, we show that mammalian spindles locally dissipate sustained force by remodeling k-fiber dynamics and structure through different physical mechanisms over time (Fig. 5). In principle, this can allow the spindle to preserve robust connections to chromosomes, and maintain its structure under force throughout mitosis. Local dissipation of force limits its impact on the rest of the spindle, providing local isolation. In turn, the timescale of such dissipation limits the timescale of mechanical memory in the spindle. By regulating force dissipation, the spindle could set the impact force has on its structure over time to allow it to respond to different mechanical cues and perform different mechanical functions. Looking forward, it will be of interest to map how spindles with different k-fiber dynamics and structures across species dissipate and transmit force, and thereby preserve their structural homeostasis (Nicklas and Staehly, 1967; Shimamoto and Kapoor, 2012; Itabashi et al., 2009; Crowder et al., 2015; Takagi et al., 2019). Finally, we note that the local force dissipation we observe in the spindle is a simple engineering principle by which a cellular structure may be mechanically robust, analogous to how structural engineers design sites of local force dissipation to make buildings and bridges robust to external forces.

## Acknowledgements

We thank Le Paliulis for critical microneedle manipulation advice, and Alan Verkman’s lab for the use of their microforge. We thank Alexey Khodjakov for the gift of PtK2 GFP-α-tubulin and PtK1 PA-GFP-α-tubulin cell lines and Jagesh Shah for the gift of the PtK2 EYFP-Cdc20 cell line. We thank David Agard, Maya Anjur-Dietrich, Wallace Marshall, Tim Mitchison, Dave Morgan, Dan Needleman, Adair Oesterle, Ron Vale, Orion Weiner, and members of the Fred Chang and Dumont labs for helpful discussions.

This work was supported by NIH DP2GM119177, NIH R01GM134132, NSF CAREER 1554139, the NSF Center for Cellular Construction DBI-1548297, the Rita Allen Foundation and Searle Scholars’ Program (S.D.), NSF Graduate Research Fellowships (A.F.L. and P.S.) and a UCSF Moritz-Heyman Discovery Fellowship and UCSF Lloyd Kozloff Fellowship (A.F.L.).

The authors declare no competing financial interests.

## Author Contributions

Conceptualization, A.F.L., P.S, and S.D.; Methodology, A.F.L., P.S.; Investigation, A.F.L. and P.S.; Data Curation, A.F.L. and P.S.; Software, A.F.L and P.S.; Writing – Original Draft, A.F.L.; Writing – Review and Editing, A.F.L, P.S. and S.D.; Visualization, A.F.L.; Funding Acquisition, S.D.

## Methods

### Cell culture

PtK2 cells were cultured in MEM (Invitrogen) supplemented with sodium pyruvate (Invitrogen), nonessential amino acids (Invitrogen), penicillin/streptomycin, and 10% qualified and heat-inactivated fetal bovine serum (Invitrogen) and maintained at 37°C and 5% CO_2_. PtK2 cells stably expressing human GFP-α-tubulin (gift from A. Khodjakov, Wadsworth Center) and PtK2 cells incubated with SiR-tubulin dye were both used. PtK2 cells stably expressing human EYFP-Cdc20 (gift from Jagesh Shah, Harvard Medical School) were used for Fig. 1 validation of microneedle manipulation. SiR-tubulin (Cytoskeleton, Inc.) at 100nM and 10µM verapamil (Cytoskeleton, Inc.) were incubated with cells for 45 min prior to imaging for cells not expressing GFP-tubulin. PtK1 cells stably expressing PA-GFP tubulin (gift from A. Khodjakov) were cultured in F12 media (Invitrogen) supplemented with penicillin/streptomycin, and 10% qualified and heat-inactivated fetal bovine serum (Invitrogen) and maintained at 37°C and 5% CO_2_. For photoactivation experiments, PtK1 PA-GFP tubulin cells were co-labeled with SiR-tubulin as above to mark overall spindle structure. Control cells labeled with SiR-tubulin that did not undergo microneedle manipulation still exhibited chromosome oscillations and poleward microtubule flux at a rate of 0.40 ± 0.06 µm/min (Fig. 3C), indicating that this concentration and length of dye incubation did not suppress k-fiber microtubule dynamics in these cells.

### Microscopy

Live cells were imaged using an inverted microscope (Eclipse Ti-E; Nikon) with a spinning disk confocal (CSU-X1; Yokogawa Electric Corporation), head dichroic Semrock Di01-T405/488/568/647 for multicolor imaging, equipped with 405 nm (100 mW), 488 nm (120mW), 561 nm (150mW), and 642 nm (100mW) diode lasers, emission filters ET455/50M, ET525/ 50M, ET630/75M and ET690/50M for multicolor imaging, and an iXon3 camera (Andor Technology) operated by MetaMorph (7.7.8.0; Molecular Devices). Cells were imaged with a 100x 1.45 Ph3 oil objective and 1.5x lens every 10 s acquiring 3 z-planes spaced 0.35 – 0.50 µm apart with a PZ-2000 z-piezo stage (ASI). Cells were imaged in a stage-top incubation chamber (Tokai Hit) with the top lid removed and maintained at 30°C. Cells were plated on glass-bottom 35mm dishes coated with poly-D-lysine (MatTek Corporation) and imaged in CO_2_ independent MEM (Invitrogen) supplemented as for PtK2 cell culture as described above. Photoactivation was performed using a MicroPoint pulsed laser system (Andor) to deliver several 3-ns 20Hz pulses of 405nm light to activate PA-GFP-tubulin (Fig. 3).

### Microneedle manipulation

Microneedle manipulation was adapted for use in mammalian spindles for sustained periods of many minutes by optimizing needle dimensions, contact geometry, and speed of motion to minimize cellular damage. Further microneedle manipulation details can be found in (Suresh et al., 2019).

#### Preparation of microneedles

Glass capillaries with an inner and outer diameter of 1 mm and 0.58 mm respectively (1B100-4 and 1B100F-4, World Precision Instruments) were used to create microneedles using a pipette puller (P-87, Sutter Instruments, Novato, CA). For a ramp value of 504 (specific to the type of glass capillary and micropipette puller), we used the following settings: Heat = 509, Pull = 70, velocity = 45, delay = 90, pressure = 200, prescribed to generate microneedles of 0.2 µm outer tip diameter (Sutter Instruments Pipette Cookbook). The measured diameter of the microneedle in the z-plane of the manipulated k-fiber was 1.2 ± 0.1 µm (the tip was placed deeper than the k-fiber to ensure that it would not slip during movement). Microneedles with longer tapers and smaller tips than above were more likely to rupture the cell membrane. Microneedles were bent ∼1.5 mm away from their tip at a 45° angle using a microforge (Narishige International, Amityville, NY). This allowed for microneedles placed in the manipulator at a 45° angle to approach the cell vertically and minimize the overall surface area of contact between the microneedle and the cell membrane.

Microneedles used for manipulation were coated with BSA Alexa Fluor 647 (A-34785, Invitrogen) or 555 conjugate (A-34786, Invitrogen) by soaking in the solution for 60 s before imaging (Sasaki et al., 2012): BSA-Alexa-647 and Sodium Azide (Nacalai Tesque, Kyoto, Japan) were dissolved in 0.1 M phosphate-buffered saline at the final concentration of 0.02% and 3 mM, respectively. Tip labeling was critical towards improving cell heath during sustained manipulations because it allowed us to better visualize the microneedle tip in fluorescence along with the spindle and prevented us from going too deeply into the cell, thereby causing rupture.

#### Selection of cells

Cells for micromanipulation were chosen based on being at metaphase, being flat, with a spindle having two poles in the same focal plane. These criteria were important for pulling on single k-fibers close to the top of the cell and simultaneously being able to image the whole spindle’s response over several minutes of manipulation. Cells were included in our datasets if they did not appear negatively affected by micromanipulation. We did not include cells that underwent sudden and continuous blebbing upon microneedle contact, cells with spindles that started to collapse during manipulation or cells with decondensed chromosomes.

#### Manipulation

Manipulations were performed in 3D using an xyz stepper micromanipulator (MP-225 Sutter Instruments). A 3-knob controller (ROE-200, Sutter Instruments) connected to the manipulator and controller (MPC-200, Sutter Instruments) allowed fine manual movements and was used to find and position the microneedle before imaging. To find and position the microneedle, we first located and centered the microneedle tip in the field of view using a low magnification objective (20X 0.5NA Ph1 air). We placed the microneedle in focus just above the coverslip before switching to a 100X 1.45 Ph3 oil objective and refined the xyz position of the microneedle to be right above a cell of interest, using the Ph1 phase ring to confirm microneedle position (phase ring mismatch visually highlights the position of the glass microneedle).

Upon choosing a cell to manipulate, we identified an outer k-fiber in a plane close to the top of the cell focused on this k-fiber. Then, we slowly brought the microneedle down into the cell using the fluorescent label of the microneedle tip to inform on its position until just deeper than the k-fiber of interest. If the microneedle’s position was too far away from the k-fiber of interest, we slowly moved the microneedle out of the cell, adjusted its xy position and brought it back down into the cell. Through this iterative process, we could correctly position the microneedle such that it was inside the spindle, next to the outer k-fiber.

Once the microneedle was positioned next to an outer k-fiber near the top of the cell, it was moved in a direction that is roughly perpendicular (∼60°-90°) from the spindle’s long axis using software (Multi-Link, Sutter Instruments). We wrote a custom program to take as inputs the desired angle, duration, and distance for the microneedle movement and then output a set of instructions in steps, x, y positions, and delays for the Multi-Link software to achieve to desired movement. For all manipulations except those in Fig. 4E, we moved the microneedle at 5.2 ± 0.2 µm/min for 3.1 ± 0.3 min (Fig. 1B). For the ‘pull and hold’ experiments, we moved the microneedle at 4.5 ± 0.7 µm/min for 1.7 ± 0.2 min and then held in place until breakage (Fig. 4E). At the end of the manipulation the microneedle was manually removed from the cell in the z-axis slowly (<5 µm /min) to avoid membrane rupture or cell detachment from the coverslip.

### Tracking of spindle features

For all analyses (Fig. 2-4), k-fibers were manually tracked in Fiji (version 2.0.0-rc-68/1.52g) (Schindelin et al., 2012) by drawing segmented lines along maximum intensity projections of three z planes of the fluorescent image with “spline fitting” checked. Splines were drawn from the edge of the tubulin signal at the plus-end to the center of the area of high tubulin intensity at the pole since we cannot determine specifically the location of the minus-end of the k-fiber. Spline x and y coordinates were saved in CSV files using a custom macro in Fiji and imported into Python. All subsequent analysis and plotting was performed in Python. Microneedle position was tracked using the mTrackJ plugin (Meijering et al., 2012) in Fiji using the “snap to bright centroid” feature and coordinates were saved in CSV files and imported into Python for further analysis.

### Quantification of spindle features

Pole and kinetochore position were calculated using the x and y coordinates of the point at the end of the spline that terminated at the pole and kinetochore, respectively. Time t = 0 was set to the first frame after the start of microneedle movement. Pole, microneedle, and kinetochore speed were calculated using the average displacement of the ends of the spline or center of the microneedle position over time (Fig. 2D,H). K-fiber length and net growth rate were calculated using the length of the spline over time and with linear regression from the start and end of the manipulation (Fig. 2E-I). For the analysis of k-fiber growth rate of unmanipulated k-fibers specifically during the growth phase (Fig. 2G), the start and end points were selected manually when there were at least three consecutive timepoints where the k-fiber length increased. The distance between the microneedle and plus-end was calculated as the linear distance between the center of the microneedle centroid and the plus-end terminus of the spline (Fig. 2I). Microtubule ‘splaying’ was manually scored as the first frame in which new microtubule density appeared on the side of the k-fiber near the point of high curvature (Fig. 4B,F). These events occurred within one time point (<10s), thus their dynamics of appearance could not be carefully characterized under these imaging conditions. K-fiber breakage was manually scored as the first frame in which the two k-fiber pieces moved in an uncorrelated manner (Fig. 4B,E-G).

### Photomark analysis

For photomark analysis, splines were tracked on maximum intensity projections of three z-planes using the 647 channel (SiR-tubulin label) and then that spline with a thickness of 5 pixels was used to calculate the intensity in the 488 channel (PA-GFP tubulin) at each point using a custom-written macro in Fiji, with all subsequent analysis in Python. Photomark position over time was calculated using the position along the curved k-fiber spline at which the maximum intensity value occurred after masking bright intensity directly at the pole that was separate from the photomark signal (Fig. 3C-E). Points were only included if the photomark remained in focus above background fluorescence. K-fiber intensity was normalized to the average intensity of the k-fiber in the timepoint prior to photomarking to identify the peak, however no intensity measurements were performed due to fluctuation of the k-fiber in the z-axis beyond the 3 z-planes measured. For calculation of photomark width (Fig. 3F), Gaussian fitting was performed on the normalized k-fiber intensities and the full-width at the half-maximal intensity (FWHM) was calculated using the width of the distribution (σ) obtained from the fit, as per FWHM 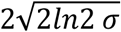 (Fig. 3G) for the subset of timepoints where the Gaussian function could fit the data.

### Curvature analysis

For curvature analysis (Fig. 4C,D), local radius of curvature (µm) was calculated by inscribing a circle through three points spaced by an interval of 1.5 µm along the spline using a custom Python script. This radius was used to calculate curvature (1/µm) by taking the inverse.

### Video preparation

Videos were formatted for publication using Fiji and set to play at 15 frames per second.

### Statistical analysis

Data are reported as mean ± SEM where indicated. All statistical testing was performed using the Python SciPy statistical package in Python. Two-sided Mann-Whitney U testing was used to compare independent samples while Wilcoxon signed-rank tests were used to compare paired data sets since we did not test whether assumptions for normality were met due to low sample size. Correlations were examined by calculating the Spearman rank-order correlation coefficient and no outliers were removed. Due to the technical challenges of these experiments, sample sizes are small. We used p < 0.05 as the threshold for statistical significance and have directly indicated in the figure and figure legend the p value and n, where n refers to the number of cells. We have therefore not performed statistical analysis for experiments with n ≤ 4 (Fig. 3). No statistical methods were used to predetermine sample size. The experiments were not randomized.

---

**Video 1: Microneedle manipulation to exert sustained force on the mammalian k-fiber, Related to Figure 1**

Microneedle manipulation of individual k-fiber in metaphase PtK2 cell to probe how k-fibers dynamics and structure respond to sustained force. The microneedle (Alexa-555, yellow) exerts force for minutes and moves the spindle (kinetochores, Cdc20-YFP, green; tubulin, SiR-tubulin, magenta) and deforms k-fibers. Manipulated spindles typically progress to anaphase (here at 10:04). Scale bar = 4 µm. Time in min:sec. Video was collected using a spinning disk confocal microscope at 1 frame every 4 s. Movie has been adjusted to play back at a constant rate of 15 frames per second. Movie corresponds to still images from Fig. 1D.

**Video 2: K-fibers persistently lengthen under applied force, Related to Figure 2**

Microneedle manipulation of individual k-fiber in metaphase PtK2 cell results in k-fiber (SiR-tubulin, white) lengthening and spindle translation and rotation in response to force. The microneedle (Alexa-555, yellow) exerts force for minutes starting at t = 0. Scale bar = 4 µm. Time in min:sec. Video was collected using a spinning disk confocal microscope at 1 frame every 10 s. Movie has been adjusted to play back at a constant rate of 15 frames per second. Movie corresponds to still images from Fig. 2B.

**Video 3: K-fiber lengthening under sustained force occurs by suppressing depolymerization at plus and minus-ends, Related to Figure 3**

Microneedle manipulation of individual k-fiber photomarked with PA-GFP-tubulin (white) in metaphase PtK1 cell reveals the mechanism of k-fiber lengthening under force. The microneedle (Alexa-555, yellow) exerts force on the k-fiber (SiR-tubulin, magenta) for minutes and the photomark remains a constant distance from the pole but a persistently increasing distance from the plus-end as the k-fiber lengthens, indicating a suppression of depolymerization at both ends. Scale bar = 4 µm. Time in min:sec. Video was collected using a spinning disk confocal microscope at 1 frame every 10 s. Movie has been adjusted to play back at a constant rate of 15 frames per second. Movie corresponds to still images from Fig. 3B.

**Video 4: K-fiber breakage occurs under sustained force for minutes, Related to Figure 4**

Microneedle manipulation of individual k-fiber for minutes reveals k-fiber breakage instead of detachment from the kinetochore or pole. The microneedle (Alexa-555, yellow) exerts force on the k-fiber (GFP-tubulin, white) for minutes and the k-fiber bends, lengthens, and ultimately breaks in two. Scale bar = 4 µm. Time in min:sec. Video was collected using a spinning disk confocal microscope at 1 frame every 10 s. Movie has been adjusted to play back at a constant rate of 15 frames per second. Movie corresponds to still images from Fig. 4B.

**Video 5: New k-fiber plus-ends can be stabilized after k-fiber breakage, Related to Figure 4**

Microneedle manipulation of individual k-fiber reveals an example of stabilized bundle plus-ends after k-fiber breakage. The microneedle (Alexa-555, yellow) exerts force on the k-fiber (SiR-tubulin, white) for minutes and is removed after the k-fiber breaks (purple arrowheads). The new plus-end fragment of the bundle persists for minutes while the fragment attached to the kinetochore is reincorporated into the spindle. This video shows the later timepoints and response of the cell from Video 3 where t = 0 is the start of microneedle manipulation. Scale bar = 4 µm. Time in min:sec. Video was collected using a spinning disk confocal microscope at 1 frame every 10 s. Movie has been adjusted to play back at a constant rate of 15 frames per second. Movie corresponds to still images from Fig. 4G.

